# Conserving Phylogenetic Diversity can be a Poor Strategy for Conserving Functional Diversity

**DOI:** 10.1101/137521

**Authors:** Florent Mazel, Arne Mooers, Giulio Valentino Dalla Riva, Matthew W. Pennell

## Abstract

For decades, academic biologists have advocated for making conservation decisions in light of evolutionary history. Specifically, they suggest that policymakers should prioritize conserving phylogenetically diverse assemblages. The most prominent argument is that conserving phylogenetic diversity (PD) will also conserve diversity in traits and features (functional diversity; FD), which may be valuable for a number of reasons. The claim that PD-maximized (‘maxPD’) sets of taxa will also have high FD is often taken at face value and in cases where researchers have actually tested it, they have done so by measuring the phylogenetic signal in ecologically important functional traits. The rationale is that if traits closely mirror phylogeny, then saving the maxPD set of taxa will tend to maximize FD and if traits do not have phylogenetic structure, then saving the maxPD set of taxa will be no better at capturing FD than criteria that ignore PD. Here, we suggest that measuring the phylogenetic signal in traits is uninformative for evaluating the effectiveness of using PD in conservation. We evolve traits under several different models and, for the first time, directly compare the FD of a set of taxa that maximize PD to the FD of a random set of the same size. Under many common models of trait evolution and tree shapes, conserving the maxPD set of taxa will conserve more FD than conserving a random set of the same size. However, this result cannot be generalized to other classes of models. We find that under biologically plausible scenarios, using PD to select species can actually lead to less FD compared to a random set. Critically, this can occur even when there is phylogenetic signal in the traits. Predicting exactly when we expect using PD to be a good strategy for conserving FD is challenging, as it depends on complex interactions between tree shape and the assumptions of the evolutionary model. Nonetheless, if our goal is to maintain trait diversity, the fact that conserving taxa based on PD will not reliably conserve at least as much FD as choosing randomly raises serious concerns about the general utility of PD in conservation.

---

In the face of the current biodiversity crisis, society needs to decide how to distribute limited funds and effort to conservation. Conservation biologists and policy makers have presented many proposals for making rational and scientific decisions about which species warrant the most protection (Bottrill et al. 2008).

One prominent prioritization scheme uses evolutionary history to place a quantitative value on species and sets of species. The idea is that when making conservation policy, we should try to conserve the set of species or habitats that harbour the greatest amount of evolutionary history (Vane-Wright et al. 1991). While there are many, overlapping metrics for measuring the evolutionary history encompassed by a set of species (Winter et al. 2013; Tucker et al. 2016), the most common is the sum of all branch lengths connecting a set of species to a common root (Faith 1992), called Phylogenetic Diversity (PD). This measure is vague insofar as the units of “branch length” are unspecified, but it is the metric whose maximization has been proposed as a conservation prioritization strategy.

While PD has been only sparingly used in actual policy decisions (for one example, see the EDGE program of the Zoological Society of London; www.edgeofexistence.org), it has caught the attention of researchers; according to Google Scholar, the original Faith (1992) paper on the topic has been cited more than 1900 times as of April 2017. Indeed, Faith’s paper has spawned an entire subfield in which biologists and mathematicians have worked out complex solutions to measuring and maximizing PD (e.g. Rodrigues and Gaston 2002; Forest et al. 2007; Bordewich et al. 2008; Bennett et al. 2014; Chao et al. 2015; Pollock et al. 2015; Thuiller et al. 2015). Faith and other researchers have proposed several key reasons why conserving PD is worthwhile. Prioritizing species’ conservation to maximize PD may help ensure that: (i) remarkable species that occur as evolutionary isolated lineages (e.g. tailed frogs, tuataras, *Welwitschia*) are prioritized (Rosauer and Mooers 2013); (ii) essential ecosystem functions and services are maintained (Cadotte et al. 2008, but see Srivastava and Vellend, 2005 for a discussion on the link with applied conservation); and that (iii) we maximize ‘evolutionary potential’ (Faith 1992; Forest et al. 2007). All of these ideas have are underpinned by the claim that phylogenetically diverse sets of taxa contain a disproportionately large amount of trait/feature/functional diversity. Hereafter, we will not make a distinction between trait, feature and functional diversity and we will refer to them as functional diversity, or FD.

Like evolutionary history, functional diversity is an ambiguous concept with many potential measures. Villéger et al. (2008) suggest that FD has three components: richness, divergence, and evenness. Functional richness generally measures ‘how much trait space is filled, while functional divergence and evenness indices describe how this space is filled’ (Schleuter et al. 2010). Functional richness represents the amount of functional trait space that is encapsulated by a set of species, is usually correlated with species richness, and can be related to the functioning of ecosystems (see, e.g. Cadotte et al. 2011). The second component, functional divergence, is largely independent of species richness and describes how species are clustered in trait space, which may be valuable to conservation biologists interested in, e.g., ecosystem services (e.g. Díaz et al. 2007). These two classes of measure are often used in trait-ecology and conservation (e.g. Devictor et al. 2010; Mouillot et al. 2014) and we assume here that conserving functional richness and/or divergence is a valuable conservation objective. While functional divergence relates to some measure of mean trait distances between species, functional evenness relates to the variance of these trait distances. This last FD dimension describes the extent to which species are clustered with their (direct) neighbours versus being regularly spaced in trait space. We did not consider any measure of functional evenness (such as the Functional Evenness Index, Villéger et al. 2008) in what follows because could not identify any potential causal link between evenness in trait space and ecosystem function or services, and trait evenness has generally not been a concern of conservation biologists.

In this paper, we ask whether maximizing PD help to conserve functional diversity. The common rationale for using PD as a proxy for FD is that many ecologically relevant traits harbour some degree of phylogenetic signal (see, e.g., Winter et al. 2013). At a glance, this seems logical: if the data shows strong phylogenetic signal, then picking distantly related taxa seems a sensible way to ensure that you have captured species from across trait space. And indeed, if we assume that traits have evolved according to a Brownian motion (BM, Felsenstein 1985) process, then this will be true (see below). The converse is also true: if traits do not show phylogenetic signal, other methods for capturing FD are needed (see, e.g., Faith 2015). A number of studies from across evolutionary biology, ecology, and conservation biology have evaluated the amount of phylogenetic signal (measured in a variety of ways, see Münkemüller et al. 2012) in ecologically important traits (see, e.g., Freckleton et al. 2002; Blomberg et al. 2003; Chamberlain et al. 2012). Recently, Kelly et al. (2014) specifically focused on the implications of phylogenetic signal for the use of PD in conservation. They constructed trees using a wide variety of morphological traits and found that while closely related species often shared many trait combinations, these traits were not informative for deeper splits in the tree. They argued that this was evidence that maximizing PD would not reliably maximize feature diversity.

The results of these studies (along with, likely many more) have been widely variable: some traits in some taxa in some regions contain a lot of phylogenetic signal while others do not. This led Winter et al. (2013, p 201) to conclude: “If the conservation goal is to conserve functional diversity, considering phylogenetic diversity might be either well suited or totally misleading”. We argue that there is an important and underappreciated assumption in this line of reasoning: that the degree of phylogenetic signal in some key trait(s) is indicative of the effectiveness of using PD to conserve FD.

There are two reasons to be suspicious of this assumption. First, our thinking about phylogenetic signal has been informed by considering a few simple models of trait evolution; other, completely different, classes of models may generate variation in phylogenetic signal that are far less intuitive. Second, the motivating idea is that policy makers should use PD to pick sets of taxa to conserve. These sets are, by definition, non-random and therefore may have different statistical properties from the clade as a whole. In this paper, we simulate data under different models of evolution and, for the first time to our knowledge, directly test how much FD the set of taxa that maximize PD (‘the maxPD set’ hereafter) contains compared to alternative possible sets.

Specifically, we contrast the outcome for FD conservation of conserving the maxPD set of taxa — and letting everything else go extinct — with conserving a random set of taxa of the same size. Here, random simply means conservation decisions that ignore phylogenetic position and the functional traits we are considering. As such, random sets provide a natural point of comparison to understand the properties of maxPD. We note that we are not testing whether conserving maxPD will maximize the amount of FD it is possible to conserve. While this claim is likely what some advocates of PD have in mind, and is what Kelly et al. (2014) actually aimed to test, it is a rather high bar to meet. Indeed, it is easy to concoct scenarios in which this will not hold; if, for example, traits were so labile that there was no phylogenetic signal (i.e., the “white noise” model), then we would expect that maxPD sets would contain no more or less FD on average than any other set. It therefore seems too high a bar to expect for PD to *always* maximize FD in order to declare it useful for conserving FD. Instead, we believe we must first clear a much lower bar — does prioritizing species based on maximizing PD do better at capturing FD than prioritizing a random set?

Below, we demonstrate that both the model of trait evolution and the tree shape are relevant for deciding whether or not PD is a good strategy for conserving FD. And, more surprisingly, we show that even when there is phylogenetic signal in the sampled traits, using PD to guide conservation decisions can lead to choice outcomes for conserving FD that are worse than if we were choosing randomly. This counter-intuitive result suggests that we need to re-assess both the way in which we intuitively consider phylogenetic signal in conservation biology, and the justification for phylogenetically-based prioritization.

## Methods

We wanted to test the following conjecture under a variety of evolutionary scenarios:

If we select a set *S* of *m* taxa from a clade of size *n* such that the sum of the branch lengths connecting *S* is at least as large as that stemming from any other possible subset (i.e. PD is maximized), then *S* will contain at least as much FD on average as a randomly chosen subset of size *m*.

Four things are notable about this test. First, as stated above, we are not trying to determine whether the maxPD set will actually maximize FD (i.e., that *S* would contain at least as much FD as any other set of the same size). Second, we are interested in the expectation, or average. Evolution certainly can take interesting turns such that some sub-clades span the functional diversity of the entire group (e.g., different clades of African rift cichlids have independently evolved the same breadth of functional diversity in different lakes; Muschick et al. 2012). Or, a trait important for ecosystem functioning may also evolve only once and we would like to make sure we capture this lineage (Davies et al. 2016). Average properties are critical, however, because PD’s utility in conservation comes precisely when we don’t know the traits or functions that matter; the best we can hope for is that, on average, we expect it to perform well. Third, we do not require *S* to uniquely maximize PD. We use the greedy algorithm proposed by Bordewich et al. (2008) to find our maxPD set of species *S*. For a given tree there are likely multiple, and possibly very many, sets of with the same PD as *S*. As this number will vary across simulations and could, in some case, be very large, we have chosen to select only one set per simulation. This allows us to carry out more simulations, increasing the generality of our results. And last, we are assuming that all of the taxa we select will survive and that every other taxa in the clade will go extinct with certainty. This is, of course, unreasonable and unrealistic but is useful for the purposes of illustration (see Discussion).

### Simulations

To explore a broad range of tree shapes, we simulated trees under three different diversification models. First, we simulated trees under a Yule process (no extinction). Second, to obtain trees that were more ‘tippy’ (i.e., having more speciation events close to the present), we used a coalescent model. In both cases, we simulated trees with 32 and with 64 taxa. To obtain trees that were more unbalanced than those typically produced by the Yule or coalescent processes, we simulated trees where the speciation rate evolved as a continuous trait along the tree (Rabosky 2010; Beaulieu and O’Meara 2015). This allowed some groups within a tree to diversify faster than others, with this heterogeneity being phylogenetically clustered.

To do the latter, we used R scripts from Beaulieu (2015, modified from Rabosky 2010) and set the initial speciation rate to. 06. Each tree was subsequently pruned to n = 64 and to n = 32. We then kept the 100 first trees that encompassed a wide range of imbalance values: we kept 10 trees by bins of 0.4 imbalance value (as measured by **β**, Blum et al. 2006) from -1.6 to 2. For a point of comparison, we also used fully imbalanced (**β** = -2) and balanced trees (**β** = 10) of 32 and 64 species.

To explore a range of continuous trait evolution models, we used 1) the BM model setting the drift parameter **σ**^2^ = 1 (we did not explore multiple values of **σ**^2^ because it does not influence the phylogenetic signal of the data and thus will not impact our results); 2) the Ornstein-Uhlenbeck (OU, Hansen 1997), with **σ**^2^ = 1 and **α** = {1.4,7} corresponding to half life of. 1 and. 5 for a tree with total height rescaled to 1; and 3) the early burst (EB, Harmon et al. 2010, r = -5 and -1). For discrete traits, we used the Markov model of evolution (Pagel 1994). We used a simple Markov model with 4 character states and all transitions rates equal to. 1 or 1. Speciational models, in which trait evolution occurs (at least in part) when lineages split, were also used for both continuous and discrete traits by applying a Pagel κ transformation to the original tree (Pagel 1999). We simulated datasets with N = {1, 2, 4} independently evolving traits. As we wanted to keep the simulations simple, we did not include variations such as multi-rate BM (O’Meara et al. 2006; Eastman et al. 2011) or multi-optima OU models (Butler and Kings 2004; Ingram and Mahler 2013; Uyeda and Harmon 2014). For each set of parameters and number of trait values, we simulated 1000 datasets. In each case, we also computed the phylogenetic signal contained in the data by calculating the spearman correlation between phylogenetic and trait distance matrices (following Kelly et al., 2014) in addition to more commonly used measures such as Blomberg’s K (Blomberg et al. 2003); Blomberg’ K does not allow for the possibility of “anti-signal” in traits, wherein close relatives are more dissimilar than distantly related taxa.

### Analysis

For each dataset, we selected two sets of *m* species (*m* = {8, 16}) out of the total number of *n* species in the tree (*n =* {32, 64}) for a total of four parameter combinations. One set was chosen at random and the other was a set that maximized PD (i.e., maxPD), using the algorithm of Bordewich et al. 2008 that we implemented in R. We then computed FD for both the random and the maxPD sets. Functional richness was estimated using the convex hull volume (Cornwell et al. 2006), which measures the total volume encapsulated by all species in trait space. In a single dimension, this simply equals the range of values. This broadly used metric in ecology is set monotonic with species richness, a property generally assumed desirable in conservation whereby the addition of a new species can never decrease the metric’s value (Ricotta 2005). Functional divergence was estimated using Rao’s quadratic entropy (Rao 1982; Botta-Dukát 2005), which represents the mean trait distance between pairs of species (including the null distance of a species with itself) and is highly correlated to the trait variance across tips (de Bello et al. 2016). While this index is not set monotonic with species richness, we feel that it might be of interest to test the robustness of our results. By using functional richness and functional divergence, we are able to capture both the spread of the data in trait space as well as how clustered it is, since it is not immediately clear what quantity is most relevant for the use of PD in conservation. For discrete traits, the convex hull volume is less meaningful than for continuous traits. Therefore, we used the number of unique trait states in the set as a measure of FD for discrete traits (Petchey and Gaston 2006; Mouillot et al. 2014).

For each simulation, we then computed the relative amount of FD in the two sets using the following metric:

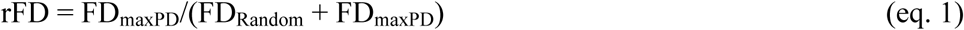

A rFD value of greater than or equal to 0.5 means that the PD set contains at least as much FD as the random set and a rFD value less than 0.5 means that it contains less. All analysis were run in R, making special use of the ape (Popescu et al. 2012), ade4 (Dray et al. 2007), phytools (Revell 2012), geiger (Pennell et al. 2014a), geometry (Habel et al. 2015), apTreeshape (Bortolussi et al. 2006), and mvMORPH (Clavel et al. 2015) packages. All code to run the analyses is available at https://github.com/FloMazel/PD_FD.

## Results

We found that, under many common models of trait evolution, conserving the maxPD set of taxa will on average conserve more FD than conserving a random set of the same size (i.e. rFD is always > .5, see Table 1, note that rFD is an average over all simulations but individual simulation may have rFD <.5). This is because related species tend to be on average closer in trait space than distantly ones (Figure 2a-d), so that selecting distantly related species increases FD. This result is more pronounced for very early evolution (as modelled by an early burst model of evolution) because in this case distantly related species are always well separated in the functional space. On the contrary, very late evolution, or very strong stabilizing selection (as modelled by the OU process) tends to erase the differences between set of species, but never leads (on average) to the maxPD set of species to harbour less FD than the random set. Overall, an increase of phylogenetic signal tends to increase the difference between FD of the two sets (Table 1 and Supp. Tables). Our results also hold for alternative tree sizes (Supp. Table 1) and Functional Divergence (measured as Rao’s Quadratic entropy, see Supp. Table 2). Also, the difference between FD of the two sets of species is largest when a small proportion of tree size is selected and tends to decrease when more species are selected (Supp. Table 2). This is expected: if 100% of the species are selected, the FD of the random and maxPD sets will be equal and equal to the FD of the entire clade (rFD = .5).

**Table 1.** For common trait macroevolution models, sets of species that maximize PD always harbour, on average, at least as much FD as random sets of species of the same size. The table presents, for each combination of macroevolutionary models (column 1), specific set of parameters (column 2-3, the transition rate for the Markov model is 1, see also methods) and number of independent traits (column 4), a measure of the relative amount of FD (rFD) between maxPD and random sets of species for pure birth Yule trees (column 5-6) and coalescent trees (column 9-10). These results correspond to a tree of 64 species from which 8 are selected either at random or to maximize PD (other combinations of these parameters are presented in Supp. Tables). The comparison of FD (as captured by the convex hull measure) between the two sets of species is quantified with the following metric: rFD=FD_maxPD_/(FD_Random_ + FD_maxPD_). A value <.5 means PD is doing worse than random, a value >.5 means PD is doing better than random and a value of. 5 means PD is doing the same as random. The phylogenetic signal for Yule trees (column 7-8) and coalescent trees (column 11-12) is measured with the Blomberg K (for multiple traits, the mean across traits is given). All statistics are based on 1000 simulations in each case.

| Evolutionary Model |  |  |  | Type of Trees |  |  |  |  |  |  |  |
| --- | --- | --- | --- | --- | --- | --- | --- | --- | --- | --- | --- |
| Type | Alpha | Beta | # traits | Yule Tree |  |  |  | Coalescent Tree |  |  |  |
|  |  |  |  | rFD |  | Phylo. Signal |  | rFD |  | Phylo. Signal |  |
|  |  |  |  | Mean | Sd | Mean | Sd | Mean | Sd | Mean | Sd |
| BM | 0 | 0 | 1 | 0.53 | 0.09 | 0.98 | 0.43 | 0.54 | 0.09 | 1.04 | 0.93 |
| BM | 0 | 0 | 2 | 0.55 | 0.13 | 1 | 0.31 | 0.6 | 0.12 | 1.01 | 0.64 |
| BM | 0 | 0 | 4 | 0.61 | 0.19 | 1 | 0.21 | 0.74 | 0.17 | 1.01 | 0.45 |
| Markov |  |  | 1 | 0.53 | 0.08 |  |  | 0.54 | 0.08 |  |  |
| Markov |  |  | 2 | 0.53 | 0.07 |  |  | 0.56 | 0.08 |  |  |
| Markov |  |  | 4 | 0.53 | 0.05 |  |  | 0.57 | 0.07 |  |  |
| EB | 0 | -5 | 1 | 0.55 | 0.09 | 4.32 | 2.51 | 0.56 | 0.11 | 9.18 | 7.36 |
| EB | 0 | -1 | 1 | 0.53 | 0.09 | 1.34 | 0.62 | 0.55 | 0.09 | 1.52 | 1.33 |
| EB | 0 | -5 | 2 | 0.62 | 0.13 | 4.33 | 1.83 | 0.66 | 0.15 | 9.32 | 5.55 |
| EB | 0 | -1 | 2 | 0.57 | 0.12 | 1.33 | 0.43 | 0.61 | 0.13 | 1.52 | 1.01 |
| EB | 0 | -5 | 4 | 0.77 | 0.16 | 4.23 | 1.36 | 0.82 | 0.16 | 9.23 | 4.17 |
| EB | 0 | -1 | 4 | 0.65 | 0.17 | 1.34 | 0.35 | 0.74 | 0.17 | 1.49 | 0.67 |
| OU | 1.4 | 0 | 1 | 0.51 | 0.09 | 0.54 | 0.16 | 0.54 | 0.09 | 0.44 | 0.3 |
| OU | 7 | 0 | 1 | 0.51 | 0.1 | 0.24 | 0.06 | 0.52 | 0.09 | 0.14 | 0.06 |
| OU | 1.4 | 0 | 2 | 0.53 | 0.13 | 0.53 | 0.12 | 0.59 | 0.13 | 0.45 | 0.22 |
| OU | 7 | 0 | 2 | 0.51 | 0.13 | 0.24 | 0.04 | 0.55 | 0.13 | 0.14 | 0.04 |
| OU | 1.4 | 0 | 4 | 0.56 | 0.2 | 0.54 | 0.08 | 0.7 | 0.17 | 0.45 | 0.16 |
| OU | 7 | 0 | 4 | 0.51 | 0.2 | 0.23 | 0.03 | 0.59 | 0.19 | 0.14 | 0.03 |

However, this result cannot be generalized to all classes of models. When traits evolve on an imbalanced tree under a speciational model (Figure 1), early diverging species are always selected to maximize PD (species No 1, 2 and 3 in Figure 2d-g) but are functionally relatively similar since their traits have not diverged much. Here, a random choice of species will, on average, select species that are much less functionally similar, yielding higher FD and thus an rFD <.5 (Figure 2d-h). As with other models, the difference between FD of the two sets of species is strongest when a small proportion of tree size is selected and tends to decrease when more species are selected (Fig. S1). This result also holds using Rao’s measure of Functional Divergence (Fig. S2).

**Figure 1.**
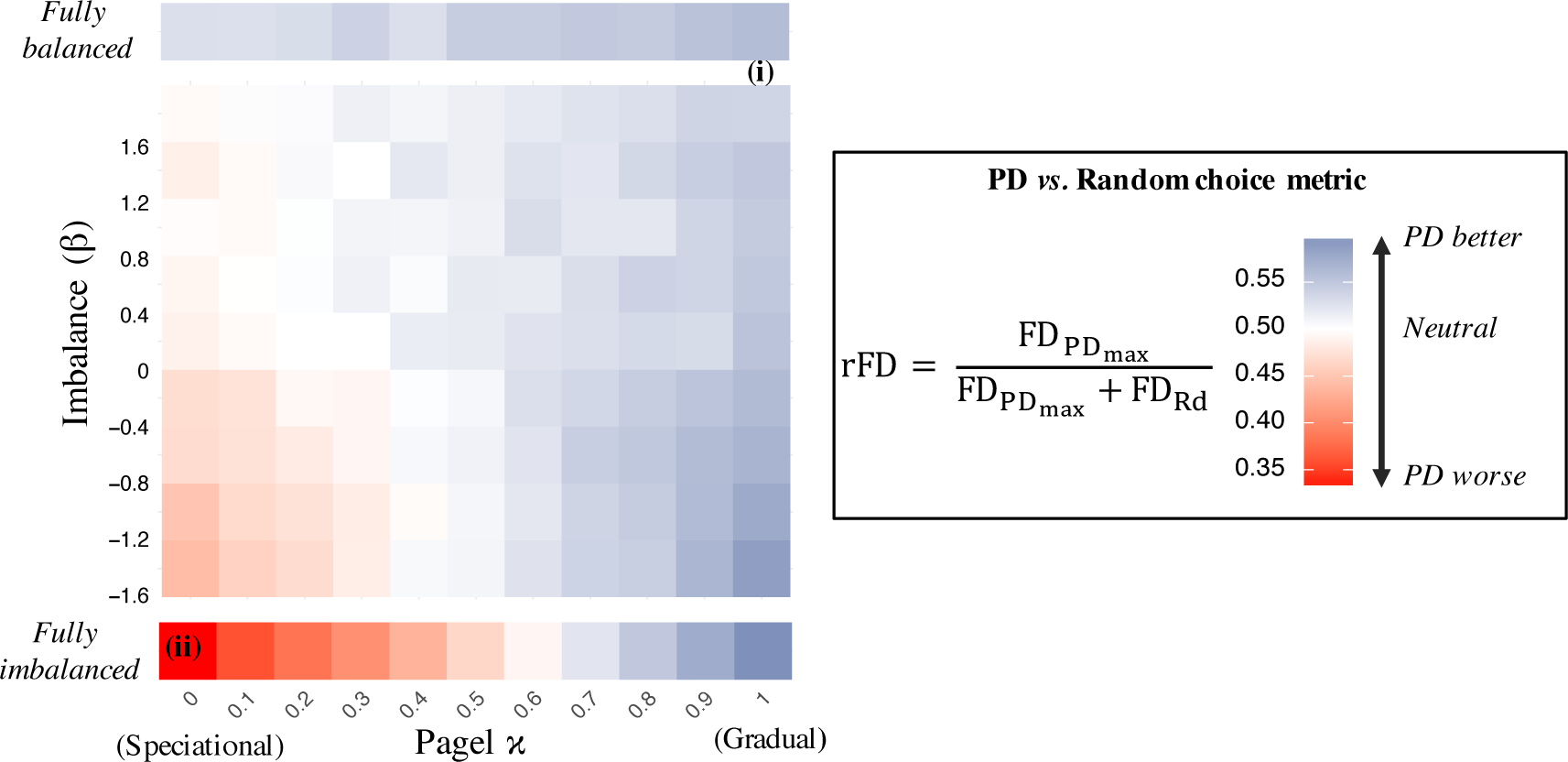
Under a speciational model trait evolution on imbalanced trees, sets of species that maximize PD harbour less FD than random sets of species of the same size. The figure represents rFD (the relative amount of FD captured by the convex hull measure between the maxPD set and random sets of species) as a function of tree imbalance (as measured by β, Y-axis) and the degree of speciational vs. gradual evolution (as measured by Pagel κ, X-axis). The color of each grid cell reflects the mean value of the metric over 100 trait simulations on 10 different trees (for a total of 1000 simulations) or, in the case of fully balanced and fully imbalanced trees, 1000 simulations on one single tree. Results are based on sets of 8 species out of 64 (tree size) and two traits. The two specific positions ‘i’ and ‘ii’ drawn on the figure refers to the parameter space position of the examples presented in Figure 2, panels a-d and e-h, respectively. The tree presented in Figure 2a (corresponding to the position marked by ‘i’ in the present figure) has an imbalance of ß=3.5.

Above we described the results for n=2 traits. Multiple traits are likely important for maintaining ecosystem functions and services and for potentially promoting diversification. However, our results do not qualitatively depend on how many traits we consider. If we use convex hull volume as a measure of FD, then the patterns we see in one or two dimensions are only exacerbated in higher trait dimensions (Fig. S1): in cases where maxPD does poorly, adding more traits makes it do worse, and in cases where it does well, more traits accentuate its success. When we measured FD using Rao’s quadratic entropy, there was no difference between results at two or higher dimensions (Fig. S2). This is because Rao’s quadratic entropy represents the mean functional distance between species (including comparing a species to itself) and we know that, for a BM model, increasing the number of traits simply decreases the variance of functional distances between species (see e.g. Letten and Cornwell 2015) and thus will not impact the average of the rFD metric. Importantly all our results are also robust to variation in tree size and number of selected species (Fig. S1-2) and also hold when a speciational model of evolution for discrete traits is applied instead (i.e. a Markov model, see Fig. S3).

After seeing our results, we naively thought that if there was a non-negative correlation between the traits and the phylogeny (i.e., “phylogenetic signal” broadly construed), this would mean that PD should on average do at least as well as random. Our intuition here was wrong. Indeed, even in our “worst case” scenario, when the tree is perfectly imbalanced and trait evolution only occurs at speciation, the correlation between the trait covariance matrix and the phylogenetic covariance matrix is still positive — close relatives resemble one another but selecting the maxPD set of taxa captures less FD than a randomly chosen set on average (Fig S4-5)! The key to resolving this apparent paradox is recognizing that the phylogenetic signal of the entire dataset is not expected to equal the phylogenetic signal of non-random subsets of the data. In particular, the set of species that maximized PD is expected to occupy a very particular position in the phylogenetic and functional distances space.

To intuitively understand this point, we present in Figure 2 (panels d-h) a simplified toy example with a fully imbalanced phylogeny of 16 species from which four species are selected, either at random (squares in the figure) or in order to maximize PD (maxPD set, represented by triangles). In this case, species 1, 2 and 3 will always be selected to maximize PD, while the fourth one will be chosen at random among the remaining species (Fig 2d). In the case of a speciational model of evolution, three out of four species from the maxPD set (species 1,2 and 3) will be, on average, relatively clumped in the trait space (Fig. 2e, triangles) and thus harbour small trait distances, while being distantly related in the phylogeny (Fig. 2g). On the contrary, the random subset (squares) will be more spread in the trait space (Fig. 2f) and thus harbour relatively higher trait distances, while being relatively less distant in the phylogeny (Fig. 2g). So, the random set will harbour more FD than the maxPD set (Fig 2h). While the overall (i.e. for all species) relationship between trait and phylogenetic distances is slightly positive (and not negative), the same relationship restricted to random and maxPD subsets becomes negative (imagine a line between squares and triangles on Figure 2g). It thus appears that the overall trend between all species are not representative of the trend between members of the maxPD and random sets; the measure of phylogenetic signal on the whole phylogeny may not be a good proxy for the representativeness of FD by the maxPD set of species.

**Figure 2.**
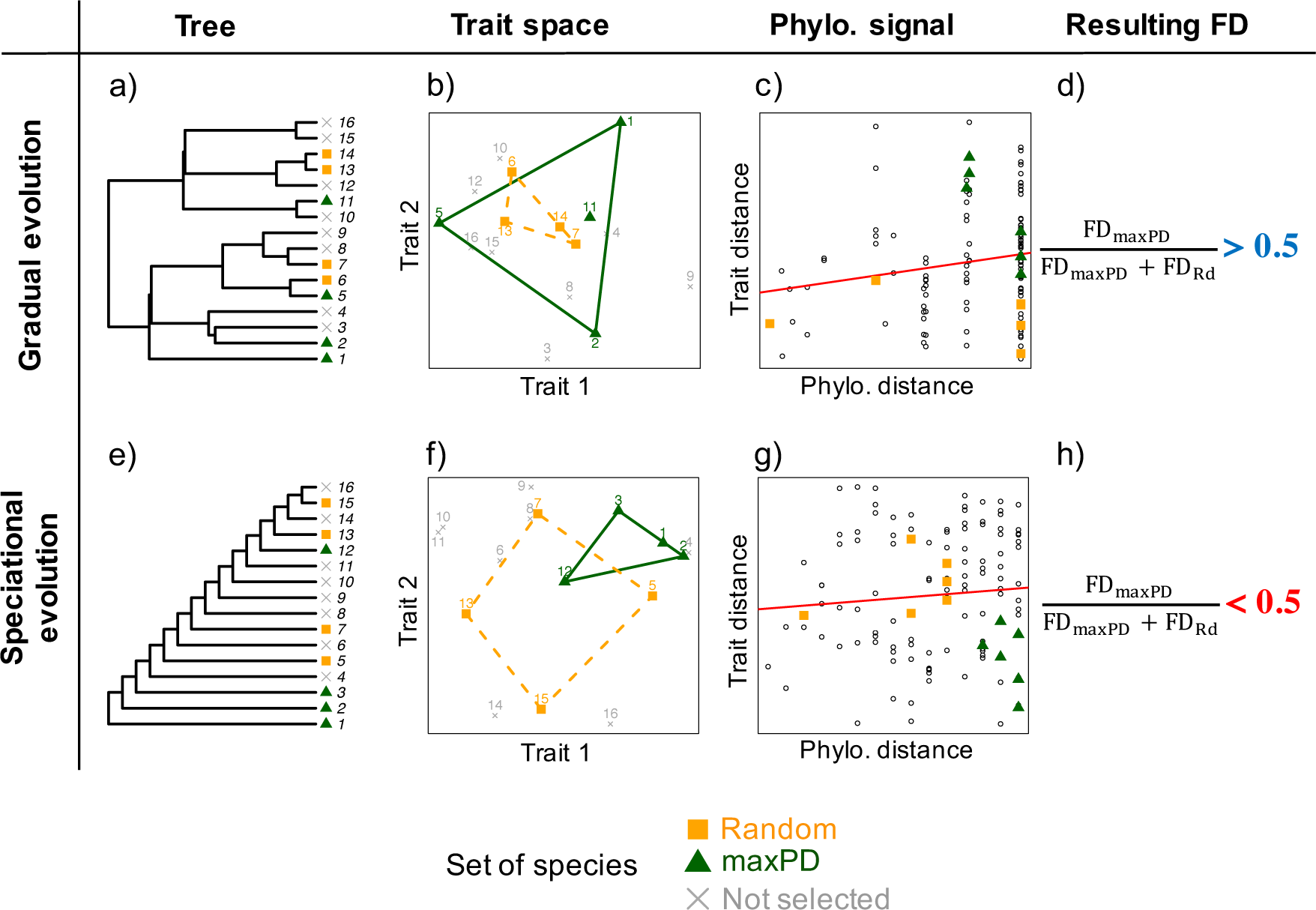
Examples of cases where the set of species that maximize PD harbours more (a-d) or less (e-h) FD than a random set of species. For each example, the original phylogenetic tree (panels a and d), the position of species in trait space and their corresponding convex hull (panels b and f), the relationship between phylogenetic and trait distances (panels c and g) and the corresponding relative amount of FD between PD_max_ set and random sets are given (panels d and h). Example (a-d) corresponds to a BM model on a relatively balanced tree while example (e-h) corresponds to a speciational model (Pagel κ = 0) on a fully imbalanced tree. Both examples are also reported in Figure 1, but note that here, for the purpose of simplicity, we used a tree with only 16 species from which four species were selected.

## Discussion

Most of the arguments for using PD in conservation decisions reason that conserving phylogenetically diverse sets of taxa is valuable because it conserves some sort of trait diversity; for other rationales for conserving PD see e.g., Vane-Wright et al. (1991) or Rosauer and Mooers (2013). Trait diversity may be valuable if it helps maintain ecosystem functioning and services (e.g. Best et al. 2013; Winter et al. 2013; Gross et al. 2017), if it captures ‘evolutionary potential’ (Faith 1992; Forest et al. 2007), or if trait diversity increases the probability of encompassing rare traits that are deemed valuable for their rareness *per se* (e.g. egg-laying in mammals, Rosauer and Mooers, 2013). Here, we are agnostic as to why traits are valuable to conserve; we only assume that they are.

Our main results speak to at least on other recent paper that also purported to test whether PD was a good proxy of feature diversity. Using a wide variety of morphological traits previously used to infer phylogenies, Kelly et al. (2014) showed that, while closely related species often share many trait combinations, these traits are not informative for deeper splits in the tree – i.e. that phylogenetic signal decays rapidly in the tested character matrices. A second key finding of the Kelly et al. study was that the trait distances between the two most distant species in the tree (i.e. considering FD of the maxPD sets of two species) is lower than the maximal trait distance in the dataset. Our test is both more stringent and more general than that of Kelly et al. First, we did not test whether preserving the maxPD species will maximize the amount of FD it is possible to preserve, but rather if the maxPD set capture more FD than a random set, a much lower bar to meet. For example, even in the situation where we found the maxPD set to harbour more FD than random (e.g. in the case of a simple BM model), it is likely that this set does not maximize FD. Second, while Kelly et al. focused on the FD of the maxPD set that comprises only two species, we consider here sets of taxa with a broader range of sizes (8 and 16 species). This allowed us to show that the measure of phylogenetic signal on the whole phylogeny may not be a good proxy for the representativeness of FD by the maxPD set of species.

Our analysis is, of course, rather oversimplified in some ways. In the real world, we do not have full control over which species survive and which are lost. Conservation prioritization itself is a result of a complex interplay of social, economic, political, and scientific priorities and is not always species-centred. And even if we did have the power to decide, we would neither conserve everything we chose, nor would everything we didn’t choose go extinct. Furthermore, the extinction proportions used in our simulations (e.g. 75%) are beyond dystopic. But the simplicity of our simulations allows us to evaluate the logic underlying the (seemingly obvious, but not actually obvious at all) claim that conserving phylogenetic diversity will result in conserving trait diversity. We realize also that some of the situations which produce rFD values of less than 0.5 may not be biologically realistic. It is unlikely that *most* trait evolution is speciational (Pennell et al. 2014b) and, while empirical trees are more unbalanced than those produced by Yule models (Mooers and Heard 1997), totally unbalanced trees are rare. While, such extreme scenarios are not necessary to reliably get rFD values of less than 0.5, we think that these cases are useful for critically evaluating the underlying logic behind the use of PD and will perhaps stimulate the production of more direct tests of the usefulness of PD to represent FD.

While there have been several meta-analyses comparing the fit of various trait models across clades (Harmon et al. 2010; Pennell et al. 2015), these have been limited to a few simple models, all of which are in the part of parameter space where PD performs well as a proxy for FD. More comprehensive meta-analyses of the fit of models to comparative data are required to allow us to assess where in model space traits of interest generally fall. Furthermore, recent innovations using simulation-based approaches (e.g. Slater et al. 2012; Sukumaran et al. 2016; Clarke et al. 2017) may allow us to expand beyond our limited set of process models. A simpler empirical test of the utility of PD is to gather empirical datasets and to repeat our analytical procedure on these. We would then be able to ask for these empirical datasets whether the maxPD set of taxa will contain more FD than a randomly chosen set. To our knowledge, no such test has been performed. While this test would not provide a definitive answer to the utility of PD, it would at least provide some indication of how concerned we should be given our results.

That said, if we had some approximate idea as to how likely it is the maxPD fails to capture FD, policy recommendations might still be difficult. If maxPD does better than random in, say, 80% of clades/traits, should this be interpreted as an endorsement of the use of PD in conservation or a denouncement? What level of increase in FD is important? A formal decision-theoretic framework (Robert 2007) might be needed for navigating these thorny problems.

## Conclusion

Given the interest in using PD in conservation decisions and the amount of work that has gone into the problem of how to measure and prioritize PD, it is surprising that there has not been direct theoretical or broad empirical evaluations of what exactly PD captures. Here, we find that under many common models of trait evolution and tree shapes, conserving the maxPD set of taxa will indeed conserve more FD than conserving a random set of the same size. However, under other biologically plausible scenarios, using PD to select species can actually lead to less FD compared to a random set. Importantly, this can occur even when there is phylogenetic signal in the traits. The fact that conserving taxa based on PD will not always reliably conserve at least as much FD as choosing randomly may raise serious concerns about the utility of PD in conservation if our goal is to save a diverse set of traits.

## Acknowledgements

This paper is a joint effort of the sCAP working group funded by the Synthesis Centre of the German Centre for Integrative Biodiversity Research Halle-Jena-Leipzig and the Canadian Institute for Ecology and Evolution. FM was supported by an NSERC Accelerator Grant to AOM. GDR thanks Mike Steel. We thank Risa Sergeant at the University of Ottawa for hospitality, the Crawford Lab for Evolutionary Studies at SFU for discussion, Luke Harmon and Will Cornwell for statistical advice, Caroline Tucker, William Pearse, Marc Cadotte, Wilfried Thuiller, Laura Pollock, Thomas Buckley, and two anonymous reviewers for comments on a previous version of this manuscript.

## Supplemental Material

### Supplemental Tables

**Supplemental Table 1.**
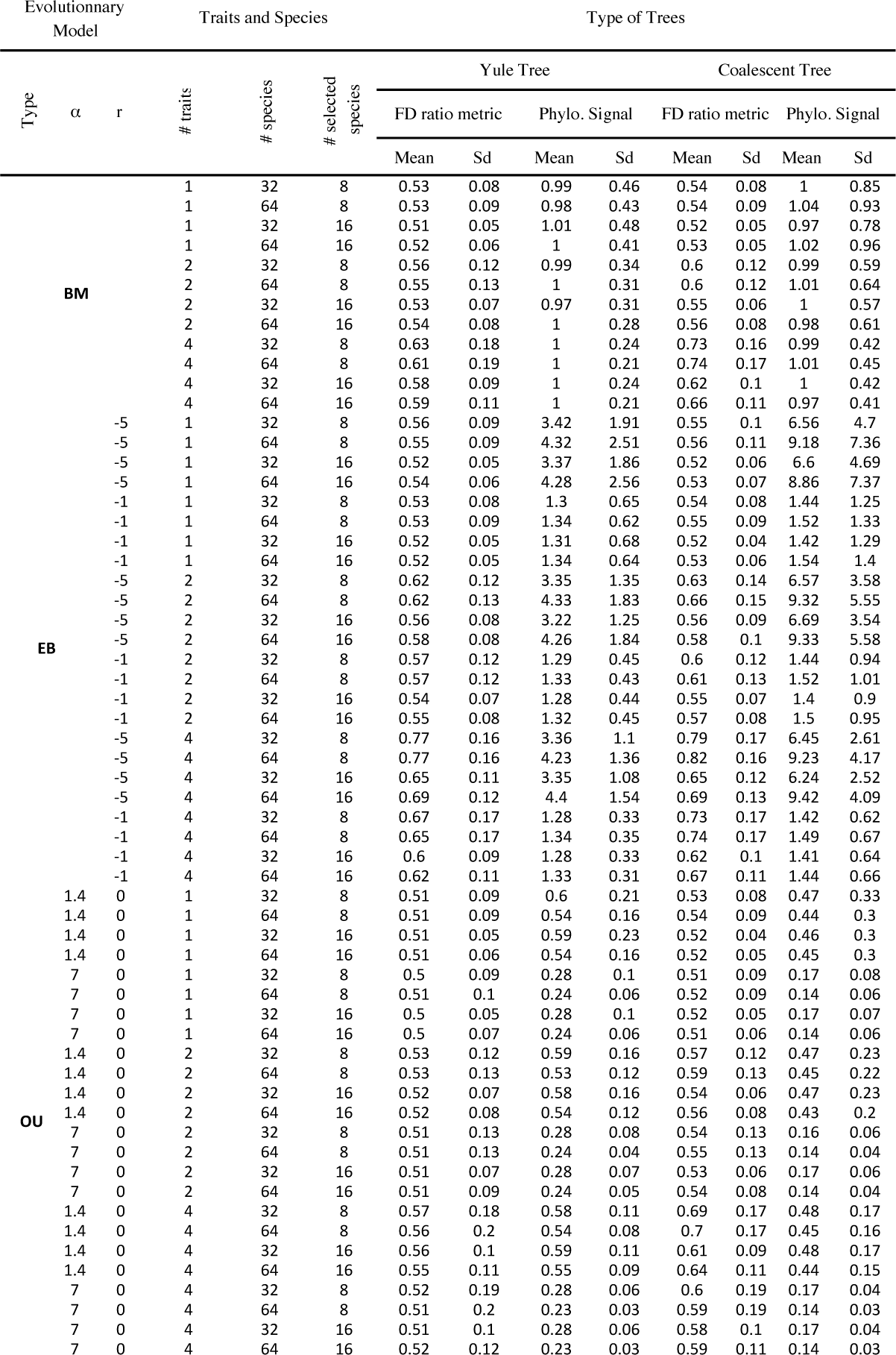
For common trait macroevolution models, sets of species that maximize PD always harbour, on average, at least as much FD (Convex hull measure) as random sets of species of the same size. The table presents, for each combination of macroevolutionnary models (column 1), specific set of parameters (column 2-3), number of independent traits (column 4), tree size (column 5) and number of selected species (column 6) a measure of the relative amount of FD between PD_max_ and random sets of species for pure birth Yule trees (column 7-8) and coalescent trees (column 11-12).The comparison of FD (<u>as measured by the convex hull measure</u>) between the two sets of species is quantified with the following metric: FD_maxPD_/(FD_Randome_ + FD_maxPD_). A value <.5 means PD is doing worse than random, a value >.5 means PD is doing better than random and a value of. 5 means PD is doing the same as random. The phylogenetic signal for Yule trees (column 9-10) and coalescent trees (column 13-14) is measured with the Bloomberg K (for multiple traits, the mean across traits is given). All statistics are based on 1000 simulations in each case.

**Supplemental Table 2.**
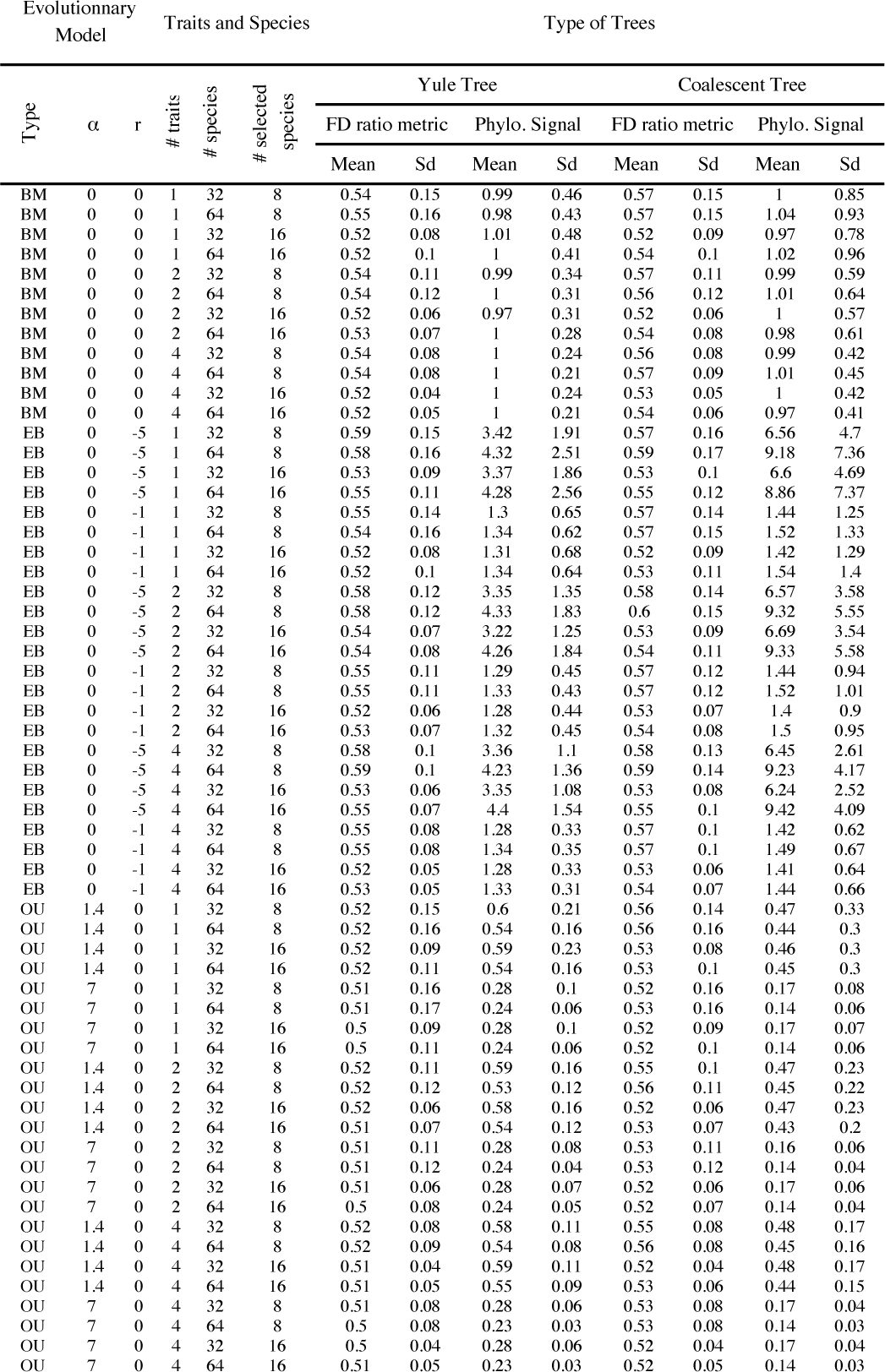
For common trait macroevolution models, sets of species that maximize PD always harbour, on average, at least as much FD (Rao quadratic entropy measure) as random sets of species of the same size. The table presents, for each combination of macroevolutionnary models (column 1), specific set of parameters (column 2-3), number of independent traits (column 4), tree size (column 5) and number of selected species (column 6) a measure of the relative amount of FD between PD_max_ and random sets of species for pure birth Yule trees (column 7-8) and coalescent trees (column 11-12).The comparison of FD (<u>as measured by the Rao quadratic entropy</u>) between the two sets of species is quantified with the following metric: FD_maxPD_/(FD_Randome_ + FD_maxPD_). A value <.5 means PD is doing worse than random, a value >.5 means PD is doing better than random and a value of. 5 means PD is doing the same as random. The phylogenetic signal for Yule trees (column 9-10) and coalescent trees (column 13-14) is measured with the Bloomberg K (for multiple traits, the mean across traits is given). All statistics are based on 1000 simulations in each case.

| Type | Evolutionary Model |  |  | Traits and Species |  |  | Type of Trees |  |  |  |  |  |  |  |
| --- | --- | --- | --- | --- | --- | --- | --- | --- | --- | --- | --- | --- | --- | --- |
| | $\alpha$ | r | # traits | # species | # selected species | | Yule Tree | | | | Coalescent Tree | | | |
|  |  |  |  |  |  |  | FD ratio metric |  | Phylo. Signal |  | FD ratio metric |  | Phylo. Signal |  |
|  |  |  |  |  |  |  | Mean | Sd | Mean | Sd | Mean | Sd | Mean | Sd |
| BM | 0 | 0 | 1 | 32 | 8 |  | 0.54 | 0.15 | 0.99 | 0.46 | 0.57 | 0.15 | 1 | 0.85 |
| BM | 0 | 0 | 1 | 64 | 8 |  | 0.55 | 0.16 | 0.98 | 0.43 | 0.57 | 0.15 | 1.04 | 0.93 |
| BM | 0 | 0 | 1 | 32 | 16 |  | 0.52 | 0.08 | 1.01 | 0.48 | 0.52 | 0.09 | 0.97 | 0.78 |
| BM | 0 | 0 | 1 | 64 | 16 |  | 0.52 | 0.1 | 1 | 0.41 | 0.54 | 0.1 | 1.02 | 0.96 |
| BM | 0 | 0 | 2 | 32 | 8 |  | 0.54 | 0.11 | 0.99 | 0.34 | 0.57 | 0.11 | 0.99 | 0.59 |
| BM | 0 | 0 | 2 | 64 | 8 |  | 0.54 | 0.12 | 1 | 0.31 | 0.56 | 0.12 | 1.01 | 0.64 |
| BM | 0 | 0 | 2 | 32 | 16 |  | 0.52 | 0.06 | 0.97 | 0.31 | 0.52 | 0.06 | 1 | 0.57 |
| BM | 0 | 0 | 2 | 64 | 16 |  | 0.53 | 0.07 | 1 | 0.28 | 0.54 | 0.08 | 0.98 | 0.61 |
| BM | 0 | 0 | 4 | 32 | 8 |  | 0.54 | 0.08 | 1 | 0.24 | 0.56 | 0.08 | 0.99 | 0.42 |
| BM | 0 | 0 | 4 | 64 | 8 |  | 0.54 | 0.08 | 1 | 0.21 | 0.57 | 0.09 | 1.01 | 0.45 |
| BM | 0 | 0 | 4 | 32 | 16 |  | 0.52 | 0.04 | 1 | 0.24 | 0.53 | 0.05 | 1 | 0.42 |
| BM | 0 | 0 | 4 | 64 | 16 |  | 0.52 | 0.05 | 1 | 0.21 | 0.54 | 0.06 | 0.97 | 0.41 |
| EB | 0 | -5 | 1 | 32 | 8 |  | 0.59 | 0.15 | 3.42 | 1.91 | 0.57 | 0.16 | 6.56 | 4.7 |
| EB | 0 | -5 | 1 | 64 | 8 |  | 0.58 | 0.16 | 4.32 | 2.51 | 0.59 | 0.17 | 9.18 | 7.36 |
| EB | 0 | -5 | 1 | 32 | 16 |  | 0.53 | 0.09 | 3.37 | 1.86 | 0.53 | 0.1 | 6.6 | 4.69 |
| EB | 0 | -5 | 1 | 64 | 16 |  | 0.55 | 0.11 | 4.28 | 2.56 | 0.55 | 0.12 | 8.86 | 7.37 |
| EB | 0 | -1 | 1 | 32 | 8 |  | 0.55 | 0.14 | 1.3 | 0.65 | 0.57 | 0.14 | 1.44 | 1.25 |
| EB | 0 | -1 | 1 | 64 | 8 |  | 0.54 | 0.16 | 1.34 | 0.62 | 0.57 | 0.15 | 1.52 | 1.33 |
| EB | 0 | -1 | 1 | 32 | 16 |  | 0.52 | 0.08 | 1.31 | 0.68 | 0.52 | 0.09 | 1.42 | 1.29 |
| EB | 0 | -1 | 1 | 64 | 16 |  | 0.52 | 0.1 | 1.34 | 0.64 | 0.53 | 0.11 | 1.54 | 1.4 |
| EB | 0 | -5 | 2 | 32 | 8 |  | 0.58 | 0.12 | 3.35 | 1.35 | 0.58 | 0.14 | 6.57 | 3.58 |
| EB | 0 | -5 | 2 | 64 | 8 |  | 0.58 | 0.12 | 4.33 | 1.83 | 0.6 | 0.15 | 9.32 | 5.55 |
| EB | 0 | -5 | 2 | 32 | 16 |  | 0.54 | 0.07 | 3.22 | 1.25 | 0.53 | 0.09 | 6.69 | 3.54 |
| EB | 0 | -5 | 2 | 64 | 16 |  | 0.54 | 0.08 | 4.26 | 1.84 | 0.54 | 0.11 | 9.33 | 5.58 |
| EB | 0 | -1 | 2 | 32 | 8 |  | 0.55 | 0.11 | 1.29 | 0.45 | 0.57 | 0.12 | 1.44 | 0.94 |
| EB | 0 | -1 | 2 | 64 | 8 |  | 0.55 | 0.11 | 1.33 | 0.43 | 0.57 | 0.12 | 1.52 | 1.01 |
| EB | 0 | -1 | 2 | 32 | 16 |  | 0.52 | 0.06 | 1.28 | 0.44 | 0.53 | 0.07 | 1.4 | 0.9 |
| EB | 0 | -1 | 2 | 64 | 16 |  | 0.53 | 0.07 | 1.32 | 0.45 | 0.54 | 0.08 | 1.5 | 0.95 |
| EB | 0 | -5 | 4 | 32 | 8 |  | 0.58 | 0.1 | 3.36 | 1.1 | 0.58 | 0.13 | 6.45 | 2.61 |
| EB | 0 | -5 | 4 | 64 | 8 |  | 0.59 | 0.1 | 4.23 | 1.36 | 0.59 | 0.14 | 9.23 | 4.17 |
| EB | 0 | -5 | 4 | 32 | 16 |  | 0.53 | 0.06 | 3.35 | 1.08 | 0.53 | 0.08 | 6.24 | 2.52 |
| EB | 0 | -5 | 4 | 64 | 16 |  | 0.55 | 0.07 | 4.4 | 1.54 | 0.55 | 0.1 | 9.42 | 4.09 |
| EB | 0 | -1 | 4 | 32 | 8 |  | 0.55 | 0.08 | 1.28 | 0.33 | 0.57 | 0.1 | 1.42 | 0.62 |
| EB | 0 | -1 | 4 | 64 | 8 |  | 0.55 | 0.08 | 1.34 | 0.35 | 0.57 | 0.1 | 1.49 | 0.67 |
| EB | 0 | -1 | 4 | 32 | 16 |  | 0.52 | 0.05 | 1.28 | 0.33 | 0.53 | 0.06 | 1.41 | 0.64 |
| EB | 0 | -1 | 4 | 64 | 16 |  | 0.53 | 0.05 | 1.33 | 0.31 | 0.54 | 0.07 | 1.44 | 0.66 |
| OU | 1.4 | 0 | 1 | 32 | 8 |  | 0.52 | 0.15 | 0.6 | 0.21 | 0.56 | 0.14 | 0.47 | 0.33 |
| OU | 1.4 | 0 | 1 | 64 | 8 |  | 0.52 | 0.16 | 0.54 | 0.16 | 0.56 | 0.16 | 0.44 | 0.3 |
| OU | 1.4 | 0 | 1 | 32 | 16 |  | 0.52 | 0.09 | 0.59 | 0.23 | 0.53 | 0.08 | 0.46 | 0.3 |
| OU | 1.4 | 0 | 1 | 64 | 16 |  | 0.52 | 0.11 | 0.54 | 0.16 | 0.53 | 0.1 | 0.45 | 0.3 |
| OU | 7 | 0 | 1 | 32 | 8 |  | 0.51 | 0.16 | 0.28 | 0.1 | 0.52 | 0.16 | 0.17 | 0.08 |
| OU | 7 | 0 | 1 | 64 | 8 |  | 0.51 | 0.17 | 0.24 | 0.06 | 0.53 | 0.16 | 0.14 | 0.06 |
| OU | 7 | 0 | 1 | 32 | 16 |  | 0.5 | 0.09 | 0.28 | 0.1 | 0.52 | 0.09 | 0.17 | 0.07 |
| OU | 7 | 0 | 1 | 64 | 16 |  | 0.5 | 0.11 | 0.24 | 0.06 | 0.52 | 0.1 | 0.14 | 0.06 |
| OU | 1.4 | 0 | 2 | 32 | 8 |  | 0.52 | 0.11 | 0.59 | 0.16 | 0.55 | 0.1 | 0.47 | 0.23 |
| OU | 1.4 | 0 | 2 | 64 | 8 |  | 0.52 | 0.12 | 0.53 | 0.12 | 0.56 | 0.11 | 0.45 | 0.22 |
| OU | 1.4 | 0 | 2 | 32 | 16 |  | 0.52 | 0.06 | 0.58 | 0.16 | 0.52 | 0.06 | 0.47 | 0.23 |
| OU | 1.4 | 0 | 2 | 64 | 16 |  | 0.51 | 0.07 | 0.54 | 0.12 | 0.53 | 0.07 | 0.43 | 0.2 |
| OU | 7 | 0 | 2 | 32 | 8 |  | 0.51 | 0.11 | 0.28 | 0.08 | 0.53 | 0.11 | 0.16 | 0.06 |
| OU | 7 | 0 | 2 | 64 | 8 |  | 0.51 | 0.12 | 0.24 | 0.04 | 0.53 | 0.12 | 0.14 | 0.04 |
| OU | 7 | 0 | 2 | 32 | 16 |  | 0.51 | 0.06 | 0.28 | 0.07 | 0.52 | 0.06 | 0.17 | 0.06 |
| OU | 7 | 0 | 2 | 64 | 16 |  | 0.5 | 0.08 | 0.24 | 0.05 | 0.52 | 0.07 | 0.14 | 0.04 |
| OU | 1.4 | 0 | 4 | 32 | 8 |  | 0.52 | 0.08 | 0.58 | 0.11 | 0.55 | 0.08 | 0.48 | 0.17 |
| OU | 1.4 | 0 | 4 | 64 | 8 |  | 0.52 | 0.09 | 0.54 | 0.08 | 0.56 | 0.08 | 0.45 | 0.16 |
| OU | 1.4 | 0 | 4 | 32 | 16 |  | 0.51 | 0.04 | 0.59 | 0.11 | 0.52 | 0.04 | 0.48 | 0.17 |
| OU | 1.4 | 0 | 4 | 64 | 16 |  | 0.51 | 0.05 | 0.55 | 0.09 | 0.53 | 0.06 | 0.44 | 0.15 |
| OU | 7 | 0 | 4 | 32 | 8 |  | 0.51 | 0.08 | 0.28 | 0.06 | 0.53 | 0.08 | 0.17 | 0.04 |
| OU | 7 | 0 | 4 | 64 | 8 |  | 0.5 | 0.08 | 0.23 | 0.03 | 0.53 | 0.08 | 0.14 | 0.03 |
| OU | 7 | 0 | 4 | 32 | 16 |  | 0.5 | 0.04 | 0.28 | 0.06 | 0.52 | 0.04 | 0.17 | 0.04 |
| OU | 7 | 0 | 4 | 64 | 16 |  | 0.51 | 0.05 | 0.23 | 0.03 | 0.52 | 0.05 | 0.14 | 0.03 |

### Supplemental Figures

**Figure S1.**
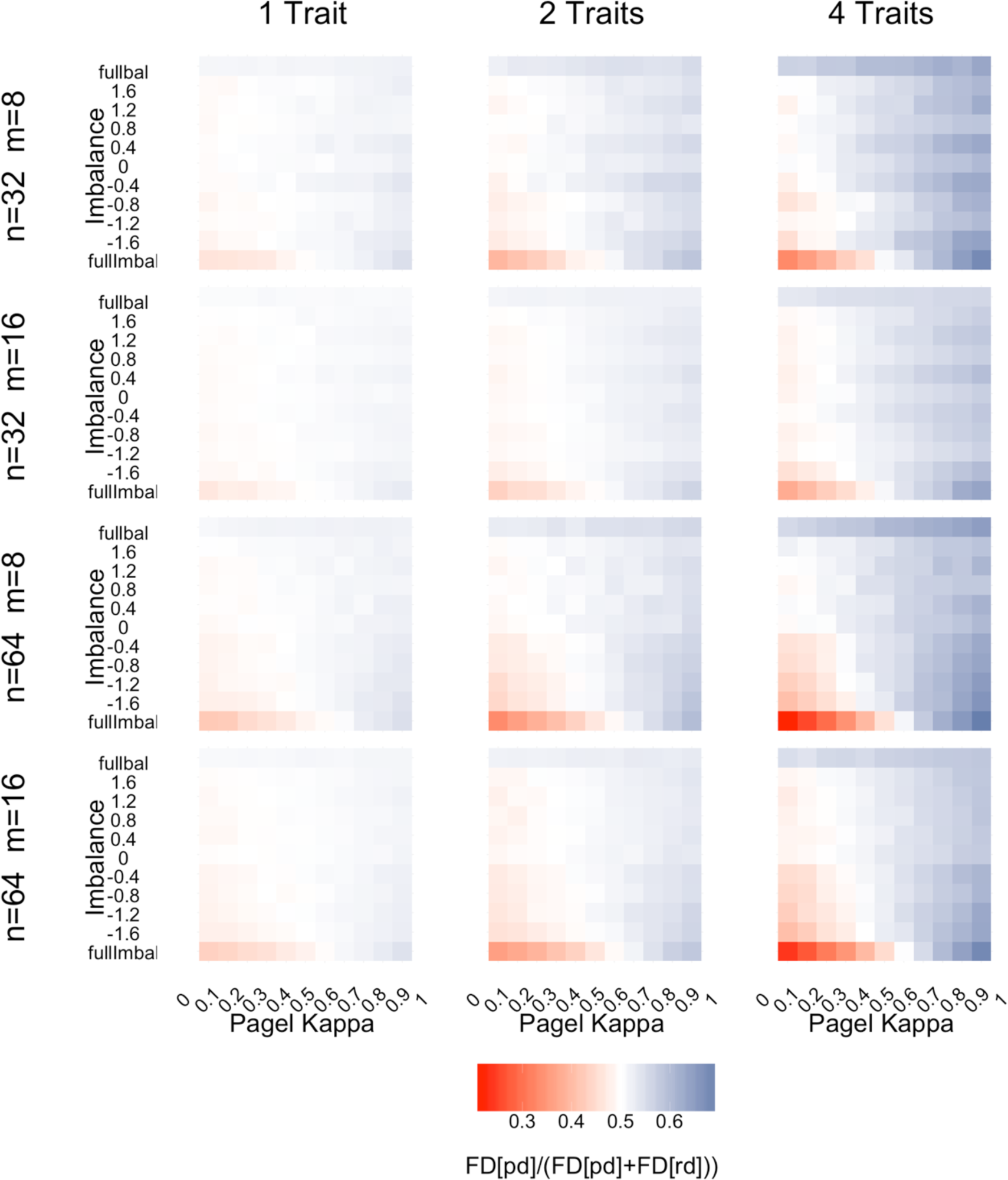
Extension of our results to multiple traits and varying tree sizes and selected species number using Convex Hull as a measure of FD. The figure present the variation of a measure of the relative amount of FD (as measured by the convex Hull measure) between PD-maximized and random set of species (see legend) in function of tree imbalance (as measured by β, Y-axis,“fullbal”refers to fully balanced tree and“fullImbal”refers to fully imbalanced tree) and the degree of speciational vs. gradual evolution (as measured by Pagel κ, X-axis). The color of each grid cell reflects the mean value of the metric over 100 trait simulations on 10 different trees (for a total of 1000 simulations) or, in the case of fully balanced and fully imbalanced trees, 1000 simulations on one single tree. Each panel corresponds to a different set of parameters (tree size (n), selected number of species (m) and number of traits).

**Figure S2.**
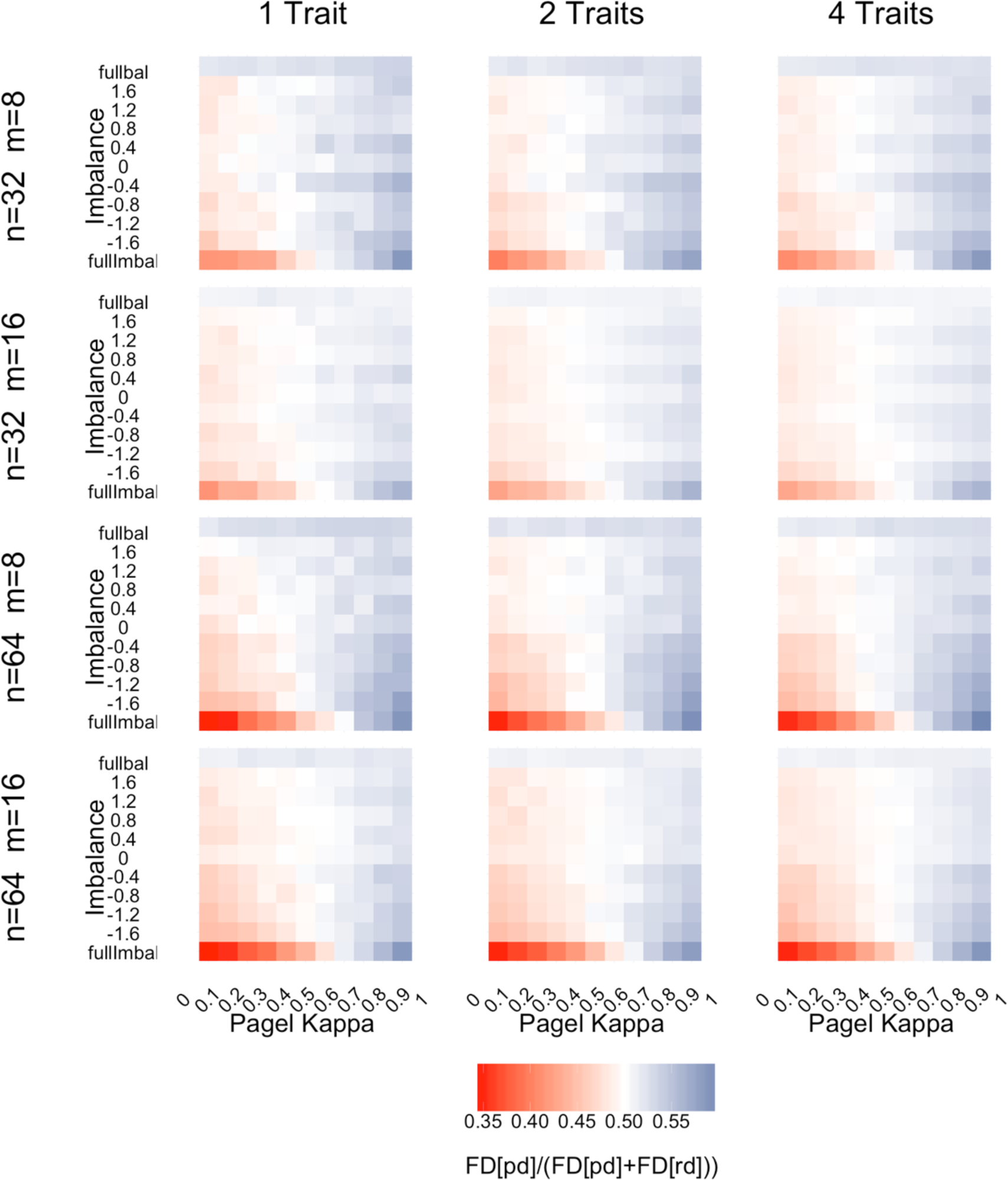
Extension of our results to multiple traits and varying tree sizes and selected species number using Rao quadratic entropy as a measure of FD. The figure presents the variation of a measure of the relative amount of FD (as measured by Rao quadratic entropy) between PD-maximized and random set of species (see legend) in function of tree imbalance (as measured by β, Y-axis,“fullbal”refers to fully balanced tree and“fullImbal”refers to fully imbalanced tree) and the degree of speciational vs. gradual evolution (as measured by Pagel κ, X-axis). The color of each grid cell reflects the mean value of the metric over 100 trait simulations on 10 different trees (for a total of 1000 simulations) or, in the case of fully balanced and fully imbalanced trees, 1000 simulations on one single tree. Each panel corresponds to a different set of parameters (tree size (n), selected number of species (m) and number of traits).

**Figure S3.**
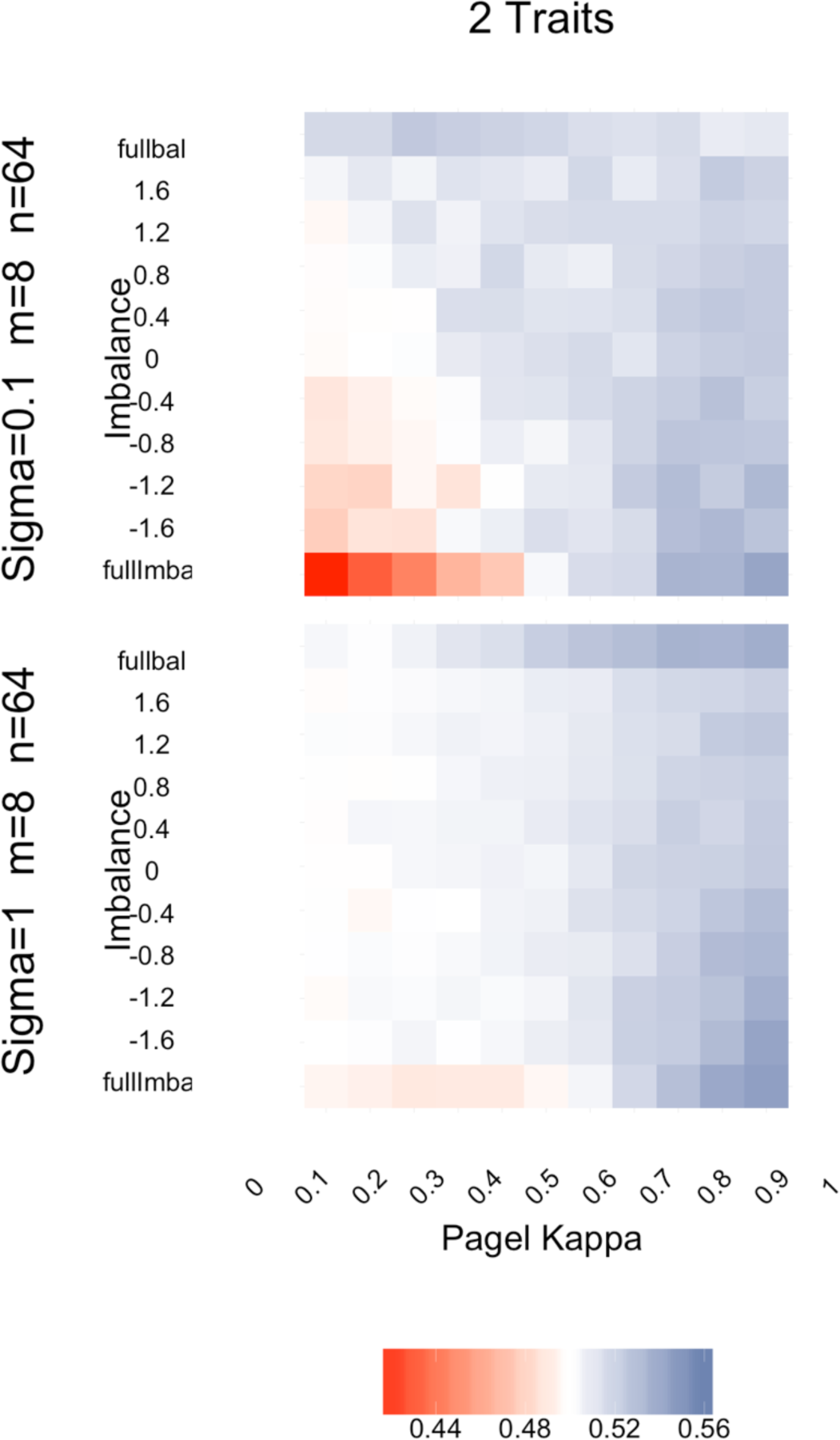
Extension of our results to discrete trait evolution. The figure presents the variation of a measure of the relative amount of FD (as measured by the number of character state combinations) between PD-maximized and random set of species (see legend) in function of tree imbalance (as measured by β, Y-axis,“fullbal”refers to fully balanced tree and“fullImbal” refers to fully imbalanced tree) and the degree of speciational vs. gradual evolution (as measured by Pagel κ, X-axis). The color of each grid cell reflects the mean value of the metric over 100 trait simulations on 10 different trees (for a total of 1000 simulations) or, in the case of fully balanced and fully imbalanced trees, 1000 simulations on one single tree. Results are based on sets of 8 species out of 64 (tree size). The different panels correspond to different values of the transition rate parameter of the Markov model.

**Figure S4.**
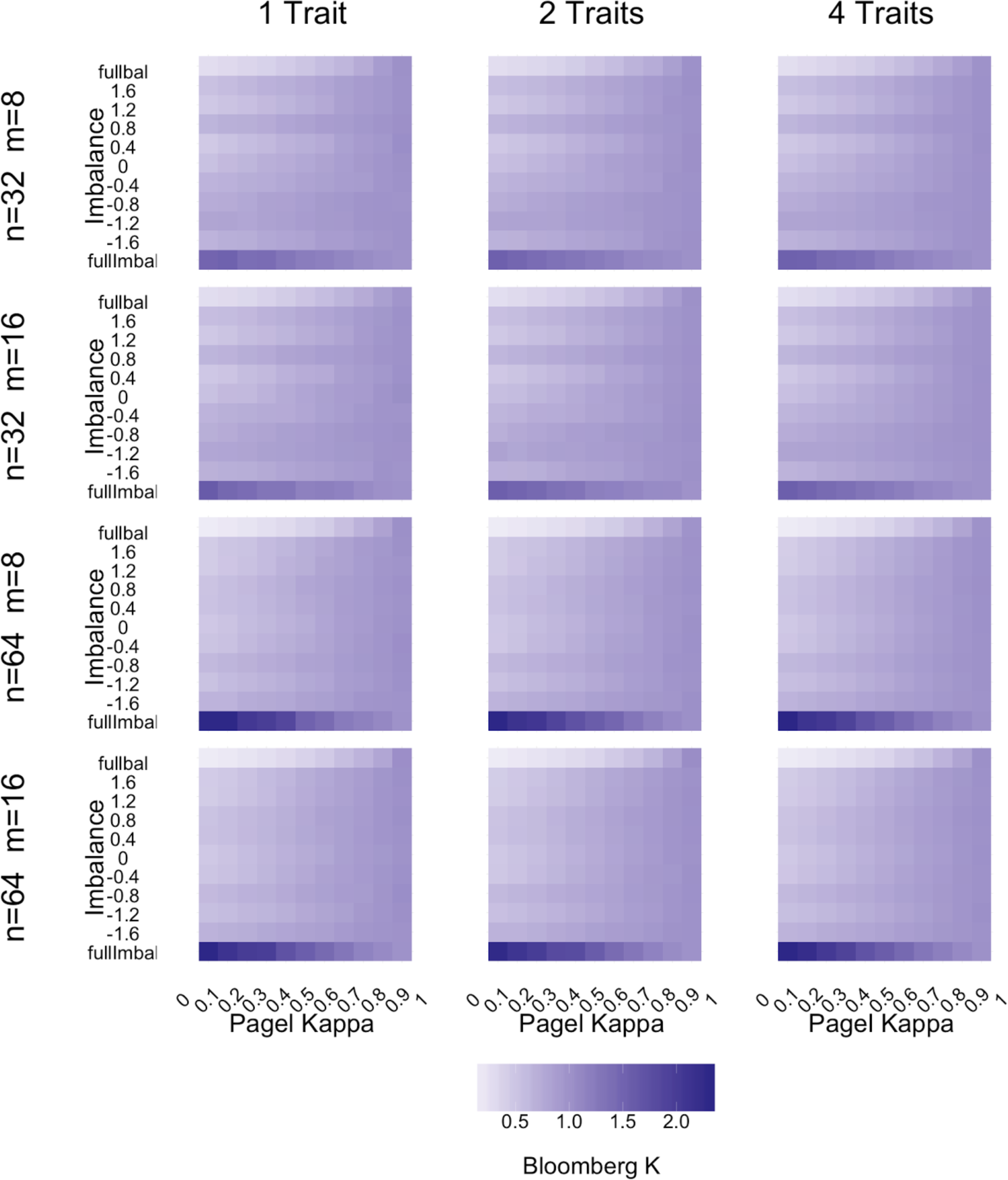
Variability of the phylogenetic signal as measured by the Bloomberg K. The figure presents the variation of phylogenetic signal across the parameter space presented on Figure 1 of the main text. Mean Bloomberg K statistic (see legend) are presented in function of tree imbalance (as measured by β, Y-axis,“fullbal”refers to fully balanced tree and“fullImbal” refers to fully imbalanced tree) and the degree of speciational vs. gradual evolution (as measured by Pagel κ, X-axis). Each panel corresponds to a different set of parameters (tree size (n), selected number of species (m) and number of traits). The color of each grid cell reflects the mean value of the metric over 100 trait simulations on 10 different trees (for a total of 1000 simulations) or, in the case of fully balanced and fully imbalanced trees, 1000 simulations on one single tree.

**Figure S5.**
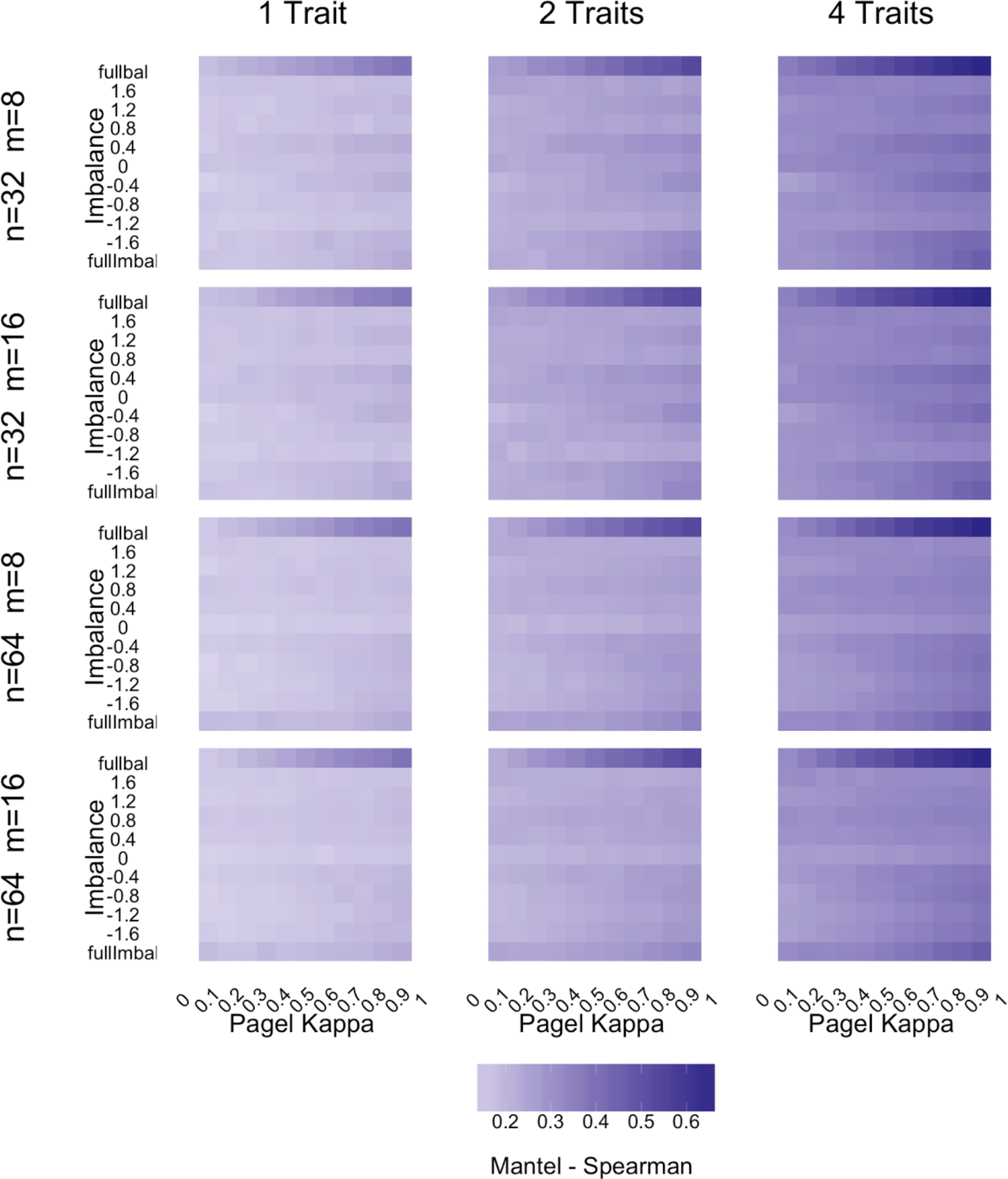
Variability of the phylogenetic signal as measured by the Spearman correlation between trait and phylogenetic distances. The figure presents the variation of phylogenetic signal across the parameter space presented on Figure 1 of the main text. Mean mantel (spearman) statistic (see legend) are presented in function of tree imbalance (as measured by β, Y-axis,“fullbal”refers to fully balanced tree and“fullImbal”refers to fully imbalanced tree) and the degree of speciational vs. gradual evolution (as measured by Pagel κ, X-axis). Each panel corresponds to a different set of parameters (tree size (n), selected number of species (m) and number of traits). The color of each grid cell reflects the mean value of the metric over 100 trait simulations on 10 different trees (for a total of 1000 simulations) or, in the case of fully balanced and fully imbalanced trees, 1000 simulations on one single tree.

## References

Beaulieu J.M., O’Meara B.C. 2015. Extinction can be estimated from moderately sized molecular phylogenies. Evolution (N. Y). 69:1036–1043.

de Bello F., Carmona C.P., Lepš J., Szava-Kovats R., Pärtel M. 2016. Functional diversity through the mean trait dissimilarity: resolving shortcomings with existing paradigms and algorithms. Oecologia. 180:933–940.

Bennett J.R., Elliott G., Mellish B., Joseph L.N., Tulloch A.I.T., Probert W.J.M., Di Fonzo M.M.I., Monks J.M., Possingham H.P., Maloney R. 2014. Balancing phylogenetic diversity and species numbers in conservation prioritization, using a case study of threatened species in New Zealand. Biol. Conserv. 174:47–54.

Best R.J., Caulk N.C., Stachowicz J.J. 2013. Trait vs. phylogenetic diversity as predictors of competition and community composition in herbivorous marine amphipods. Ecol. Lett. 16:72–80.

Blomberg S.P., Garland T.J.R., Ives A. 2003. Testing for phylogenetic signal in compartive data: behavioral traits are more labile. Evolution (N. Y). 57:717–745.

Blum M., François O., Steel M. 2006. Which Random Processes Describe the Tree of Life? A Large-Scale Study of Phylogenetic Tree Imbalance. Syst. Biol. 55:685–691.

Bordewich M., Rodrigo A., Semple C., Collins T. 2008. Selecting Taxa to Save or Sequence: Desirable Criteria and a Greedy Solution. Syst. Biol. 57:825–834.

Bortolussi N., Durand E., Blum M., Francois O. 2006. apTreeshape: statistical analysis of phylogenetic tree shape. Bioinformatics. 22:363–364.

Botta-Dukát Z. 2005. Rao’ s quadratic entropy as a measure of functional diversity based on multiple traits. J. Veg. Sci. 16:533–540.

Bottrill M.C., Joseph L.N., Carwardine J., Bode M., Cook C., Game E.T., Grantham H., Kark S., Linke S., McDonald-Madden E., Pressey R.L., Walker S., Wilson K.A., Possingham H.P. 2008. Is conservation triage just smart decision making? Trends Ecol. Evol. 23:649–654.

Butler M.A., Kings A.A. 2004. Phylogenetic comparative analysis: A modeling approach for adaptive evolution. Am. Nat. 164:683–695.

Cadotte M.W., Cardinale B.J., Oakley T.H. 2008. Evolutionary history and the effect of biodiversity on plant productivity. Proc. Natl. Acad. Sci. U. S. A. 105:17012–7.

Cadotte M.W., Carscadden K., Mirotchnick N. 2011. Beyond species: functional diversity and the maintenance of ecological processes and services. J. Appl. Ecol. 48:1079–1087.

Chamberlain S.A., Hovick S.M., Dibble C.J., Rasmussen N.L., Van Allen B.G., Maitner B.S., Ahern J.R., Bell-Dereske L.P., Roy C.L., Meza-Lopez M., Carrillo J., Siemann E., Lajeunesse M.J., Whitney K.D. 2012. Does phylogeny matter? Assessing the impact of phylogenetic information in ecological meta-analysis. Ecol. Lett. 15:627– 636.

Chao A., Chiu C.-H., Hsieh T.C., Davis T., Nipperess D.A., Faith D.P. 2015. Rarefaction and extrapolation of phylogenetic diversity. Methods Ecol. Evol. 6:380–388.

Clarke M., Thomas G.H., Freckleton R.P. 2017. Trait Evolution in Adaptive Radiations: Modeling and Measuring Interspecific Competition on Phylogenies. Am. Nat. 189:121–137.

Clavel J., Escarguel G., Merceron G. 2015. mv morph?: an r package for fitting multivariate evolutionary models to morphometric data. Methods Ecol. Evol. 6:1311–1319.

Cornwell W.K., Schwilk D.W., Ackerly D.D., Schwilk L.D.W. 2006. A trait-based test for habitat filtering: convex hull volume. Ecology. 87:1465–1471.

Davies T.J., Urban M.C., Rayfield B., Cadotte M.W., Peres-Neto P.R. 2016. Deconstructing the relationships between phylogenetic diversity and ecology: a case study on ecosystem functioning. Ecology. 97:2212–2222.

Devictor V., Mouillot D., Meynard C., Jiguet F., Thuiller W., Mouquet N. 2010. Spatial mismatch and congruence between taxonomic, phylogenetic and functional diversity: the need for integrative conservation strategies in a changing world. Ecol. Lett. 13:1030–1040.

Díaz S., Lavorel S., de Bello F., Quétier F., Grigulis K., Robson T.M. 2007. Incorporating plant functional diversity effects in ecosystem service assessments. Proc. Natl. Acad. Sci. U. S. A. 104:20684–9.

Dray S., Dufour A.-B., Dray S., Dufour A.-B. 2007. The ade4 Package: Implementing the Duality Diagram for Ecologists. J. Stat. Softw. 22.

Eastman J.M., Alfaro M.E., Joyce P., Hipp A.L., Harmon L.J. 2011. A novel comparative method for identifying shifts in the rate of character evolution on trees. Evolution. 65:3578–89.

Faith D.P. 1992. Conservation evaluation and phylogenetic diversity. Biol. Conserv. 61:1–10.

Faith D.P. 2015. The unimodal relationship between species’ functional traits and habitat gradients provides a family of indices supporting the conservation of functional trait diversity. Plant Ecol. 216:725–740.

Felsenstein J. 1985. Phylogenies and the comparative method. Am. Nat.:1–15.

Forest F., Grenyer R., Rouget M., Davies T.J., Cowling R.M., Faith D.P., Balmford A., Manning J.C., Procheş S., van der Bank M., Reeves G., Hedderson T.A.J., Savolainen V. 2007. Preserving the evolutionary potential of floras in biodiversity hotspots. Nature. 445:757–760.

Freckleton R.P., Harvey P.H., Pagel M. 2002. Phylogenetic analysis and comparative data: a test and review of evidence. Am. Nat. 160:712–726.

Gross N., Bagousse-Pinguet Y. Le, Liancourt P., Berdugo M., Gotelli N.J., Maestre F.T. 2017. Functional trait diversity maximizes ecosystem multifunctionality. Nat. Ecol. Evol. 1:132.

Habel K., Grasman R., Gramacy R.B., Stahel A., Sterratt D.C. 2015. geometry: Mesh Generation and Surface Tesselation. R package version 0.3-6. Available from https://cran.r-project.org/package=geometry.

Hansen T.F. 1997. Stabilizing Selection and the Comparative Analysis of Adaptation. Evolution (N. Y). 51:1341.

Harmon L.J., Losos J.B., Jonathan Davies T., Gillespie R.G., Gittleman J.L., Bryan Jennings W., Kozak K.H., McPeek M.A., Moreno-Roark F., Near T.J., Purvis A., Ricklefs R.E., Schluter D., Schulte II J.A., Seehausen O., Sidlauskas B.L., Torres-Carvajal O., Weir J.T., Mooers A. Ø. 2010. Early bursts of body size and shape evolution are rare in comparative data. Evolution (N. Y). 64:2385–2396.

Ingram T., Mahler D. 2013. SURFACE: detecting convergent evolution from comparative data by fitting Ornstein-Uhlenbeck models with stepwise Akaike Information Criterion. Methods Ecol. Evol. 4:416–425.

Kelly S., Grenyer R., Scotland R.W. 2014. Phylogenetic trees do not reliably predict feature diversity. Divers. Distrib. 20:600–612.

Letten A.D., Cornwell W.K. 2015. Trees, branches and (square) roots: why evolutionary relatedness is not linearly related to functional distance. Methods Ecol. Evol. 6:439– 444.

Mooers A.O., Heard S.B. 1997. Inferring Evolutionary Process from Phylogenetic Tree Shape. Q. Rev. Biol. 72:31–54.

Mouillot D., Villeger S., Parravicini V., Kulbicki M., Arias-Gonzalez J.E., Bender M., Chabanet P., Floeter S.R., Friedlander A., Vigliola L., Bellwood D.R. 2014. Functional over-redundancy and high functional vulnerability in global fish faunas on tropical reefs. Proc. Natl. Acad. Sci. 111:13757–13762.

Münkemüller T., Lavergne S., Bzeznik B., Dray S., Jombart T., Schiffers K., Thuiller W. 2012. How to measure and test phylogenetic signal. Methods Ecol. Evol. 3:743–756.

Muschick M., Indermaur A., Salzburger W. 2012. Convergent evolution within an adaptive radiation of cichlid fishes. Curr. Biol. 22:2362–8.

O’Meara B.C., Ané C., Sanderson M.J., Wainwright P.C. 2006. Testing for different rates of continuous trait evolution using likelihood. Evolution (N. Y). 60:922–933.

Pagel M. 1994. Detecting correlated evolution on phylogenies: a general method for the comparative analysis of discrete characters. Proc. R. Soc. London, Ser. B. 255:37– 45.

Pagel M. 1999. Inferring the historical patterns of biological evolution. Nature. 401:877– 884.

Pennell M.W., Eastman J.M., Slater G.J., Brown J.W., Uyeda J.C., FitzJohn R.G., Alfaro M.E., Harmon L.J. 2014a. geiger v2.0: an expanded suite of methods for fitting macroevolutionary models to phylogenetic trees. Bioinformatics. 30:2216–2218.

Pennell M.W., FitzJohn R.G., Cornwell W.K., Harmon L.J. 2015. Model Adequacy and the Macroevolution of Angiosperm Functional Traits. Am. Nat. 186:E33–50.

Pennell M.W., Harmon L.J., Uyeda J.C. 2014b. Is there room for punctuated equilibrium in macroevolution? Trends Ecol. Evol. 29:23–32.

Petchey O.L., Gaston K.J. 2006. Functional diversity: back to basics and looking forward. Ecol. Lett. 9:741–758.

Pollock L.J., Rosauer D.F., Thornhill A.H., Kujala H., Crisp M.D., Miller J.T., McCarthy M.A. 2015. Phylogenetic diversity meets conservation policy: small areas are key to preserving eucalypt lineages. Philos. Trans. R. Soc. Lond. B. Biol. Sci. 370:20140007.

Popescu A.-A., Huber K.T., Paradis E. 2012. ape 3.0: New tools for distance-based phylogenetics and evolutionary analysis in R. Bioinformatics. 28:1536–1537.

Rabosky D.L. 2010. Extinction rates should not be estimated from molecular phylogenies. Evolution (N. Y). 64:1816–1824.

Rao R.C. 1982. Diversity and dissimilarity coefficients: a unified approach. Theor. Popul. Biol. 21:24–43.

Revell L.J. 2012. phytools: an R package for phylogenetic comparative biology (and other things). Methods Ecol. Evol. 3:217–223.

Ricotta C. 2005. A note on functional diversity measures. Basic Appl. Ecol. 6:479–486.

Robert C.P. 2007. The Bayesian choice?: from decision-theoretic foundations to computational implementation. Springer.

Rodrigues A.S.L., Gaston K.J. 2002. Maximising phylogenetic diversity in the selection of networks of conservation areas. Biol. Conserv. 105:103–111.

Rosauer D.F., Mooers A.Ø. 2013. Nurturing the use of evolutionary diversity in nature conservation. Trends Ecol. Evol. 28:322–3.

Schleuter D., Daufresne M., Massol F., Argillier C. 2010. A user’s guide to functional diversity indices. Ecol. Monogr. 80:469–484.

Slater G.J., Harmon L.J., Wegmann D., Joyce P., Revell L.J., Alfaro M.E. 2012. Fitting models of continuous trait evolution to incompletely sampled comparative data using approximate bayesian computation. Evolution (N. Y). 66:752–762.

Srivastava D.S., Vellend M. 2005. Biodiversity-Ecosytem Function Research: Is It Relevant to Conservation? Annu. Rev. Ecol. Evol. Syst. 36:267–294.

Sukumaran J., Economo E.P., Lacey Knowles L. 2016. Machine Learning Biogeographic Processes from Biotic Patterns: A New Trait-Dependent Dispersal and Diversification Model with Model Choice By Simulation-Trained Discriminant Analysis. Syst. Biol. 65:525–545.

Thuiller W., Maiorano L., Mazel F., Guilhaumon F., Ficetola G.F., Lavergne S., Renaud J., Roquet C., Mouillot D. 2015. Conserving the functional and phylogenetic trees of life of European tetrapods. Philos. Trans. R. Soc. Lond. B. Biol. Sci. 370:20140005.

Tucker C.M., Cadotte M.W., Carvalho S.B., Davies T.J., Ferrier S., Fritz S.A., Grenyer R., Helmus M.R., Jin L.S., Mooers A.Ø., Pavoine S., Purschke O., Redding D.W., Rosauer D.F., Winter M., Mazel F. 2016. A guide to phylogenetic metrics for conservation, community ecology and macroecology. Biol. Rev. Camb. Philos. Soc.

Uyeda J.C., Harmon L.J. 2014. A Novel Bayesian Method for Inferring and Interpreting the Dynamics of Adaptive Landscapes from Phylogenetic Comparative Data. Syst. Biol. 63:902–918.

Vane-Wright R.I., Humphries C.J., Williams P.H. 1991. What to protect?—Systematics and the agony of choice. Biol. Conserv. 55:235–254.

Villéger S., Mason N., Mouillot D. 2008. New multidimensional functional diversity indices for a multifaceted framework in functional ecology. Ecology. 89:2290–2301.

Winter M., Devictor V., Schweiger O. 2013. Phylogenetic diversity and nature conservation: where are we? Trends Ecol. Evol. 28:199–204.

